# Higher inter-trial latency variability contributes to reduced visual EEG responses in schizophrenia

**DOI:** 10.64898/2025.11.28.689466

**Authors:** Dario Gordillo, Janir Ramos da Cruz, Andreas Brand, Eka Chkonia, Maya Roinishvili, Patrícia Figueiredo, Michael H Herzog

## Abstract

Patients with schizophrenia show strong impairments in visual backward masking, which are associated with reduced EEG N1 responses. However, it is currently unclear whether reduced N1 amplitudes in patients reflect attenuated neural responses or increased inter-trial latency variability, as both can lead to reduced trial-averaged responses. Previous studies using trial-averaged data cannot distinguish between these two possibilities. Here, we estimated inter-trial latency variability of the visual N1 component and found significantly increased variability in patients compared to controls. Inter-trial latency-variability was a strong predictor of the N1 amplitude in both groups. Importantly, after accounting for the effect of latency variability in the group comparison, patients continued to exhibit significantly reduced N1 amplitudes, although the effect size diminished from large to medium. These findings indicate that both higher latency variability and attenuated neural responses contribute to visual processing deficits in schizophrenia.

## Introduction

Visual dysfunctions are pathophysiological signatures of schizophrenia that contribute to perceptual and cognitive impairments^1^. Event-related potentials (ERPs) are often reduced in specific components in patients with schizophrenia compared to controls, and have been related to decreased performance in sensory and cognitive paradigms^2^. However, because conventional ERP analyses rely on averaging across trials, they provide limited insight into how inter-trial temporal fluctuations contribute to differences in behavioral and neural responses.

In classic ERP analysis, data are averaged across trials to improve signal-to-noise ratio (SNR). This approach is effective for components that exhibit a stable latency across trials. However, while early ERPs reflecting basic properties of the stimuli (e.g., onset and luminance), such as the P1, generally have a stable latency, later ERP components typically have higher inter-trial latency variability^3–5^. If inter-trial variability is large, even strong single-trial responses can appear attenuated in the average ERP.

Investigating the contribution of inter-trial variability is particularly relevant in schizophrenia, because recent studies have shown increased temporal variability of neural responses in patients^6–9^. Importantly, increased inter-trial variability in schizophrenia has been demonstrated for ERPs including the N2^10^, P50^11^, P3^9,12^, and MMN^13^, which have also been shown to be significantly reduced in patients in comparison to controls when trial-averaged data have been analyzed^2,14–16^. These results suggest that inter-trial variability might be a contributing factor to a range of deficits observed across paradigms.

The shine-through visual backward masking (VBM) paradigm has been extensively employed to study temporal dynamics of visual processing dysfunctions in schizophrenia^17–20^. In VBM, a brief target Vernier (i.e., two vertical lines with a horizontal offset) is presented, followed by a mask that interferes with target processing. Behaviorally, patients with schizophrenia show similar performance to controls when the Vernier is presented alone^21^. However, when the Vernier is masked, performance is approximately four times worse on average than that of controls^17,18^.

ERP studies of VBM have shown reduced global field power (GFP) amplitudes in patients around 200ms after stimulus onset, reflecting reduced visual N1 responses^22^. Importantly, in control conditions where only the mask is presented (i.e., there is no target), comparable N1 amplitudes between patients and controls have been observed, suggesting that reduced N1 reflects an impairment in the enhancement of the target information in patients, rather than a general deficit, such as reduced neural excitation^18,22,23^. Since temporally precise neural responses are critical for target enhancement, characterizing the N1 latency variability in VBM might offer a suitable test for impairments related to temporal imprecision in schizophrenia.

Here, we re-analyzed previously published EEG data from 142 patients with schizophrenia and 96 controls^24,25^ performing a VBM task to investigate the effect of inter-trial latency variability on the visual N1. To isolate the N1 component and estimate its latency at the single trial level, we used independent component analysis to increase signal-to-noise ratio, allowing us to quantify how much inter-trial variability contributes to the N1 amplitude differences between patients with schizophrenia and controls.

## Results

### ICA and single-trial latency estimation

Using multi-level group independent component analysis (mlGICA), we obtained a component resembling the visual N1 response (i.e., IC-N1; Figure 2A) characterized by a negative bilateral-occipital and positive frontal-central activation, and a time course with a peak at around 200ms. IC-N1 amplitudes were reduced in patients with schizophrenia in comparison to controls (Figure 1B, 2B, 3). The inverse of the ICA mixing matrix, and the coefficients from the first and second PCAs were used to obtain an individual-subject representation of this component (Figure 2C). IC-N1 peak amplitudes were defined as the minimum peak value in a 100ms window located within 125 and 275ms, which was centered at the time of the average peak latency of each condition and subject. Single-trial peak amplitudes were also obtained using two additional algorithms—template matching (i.e., Woody filter) and a graph cuts-based approach—which yielded similar results (Figure S2).

**Figure 1.**
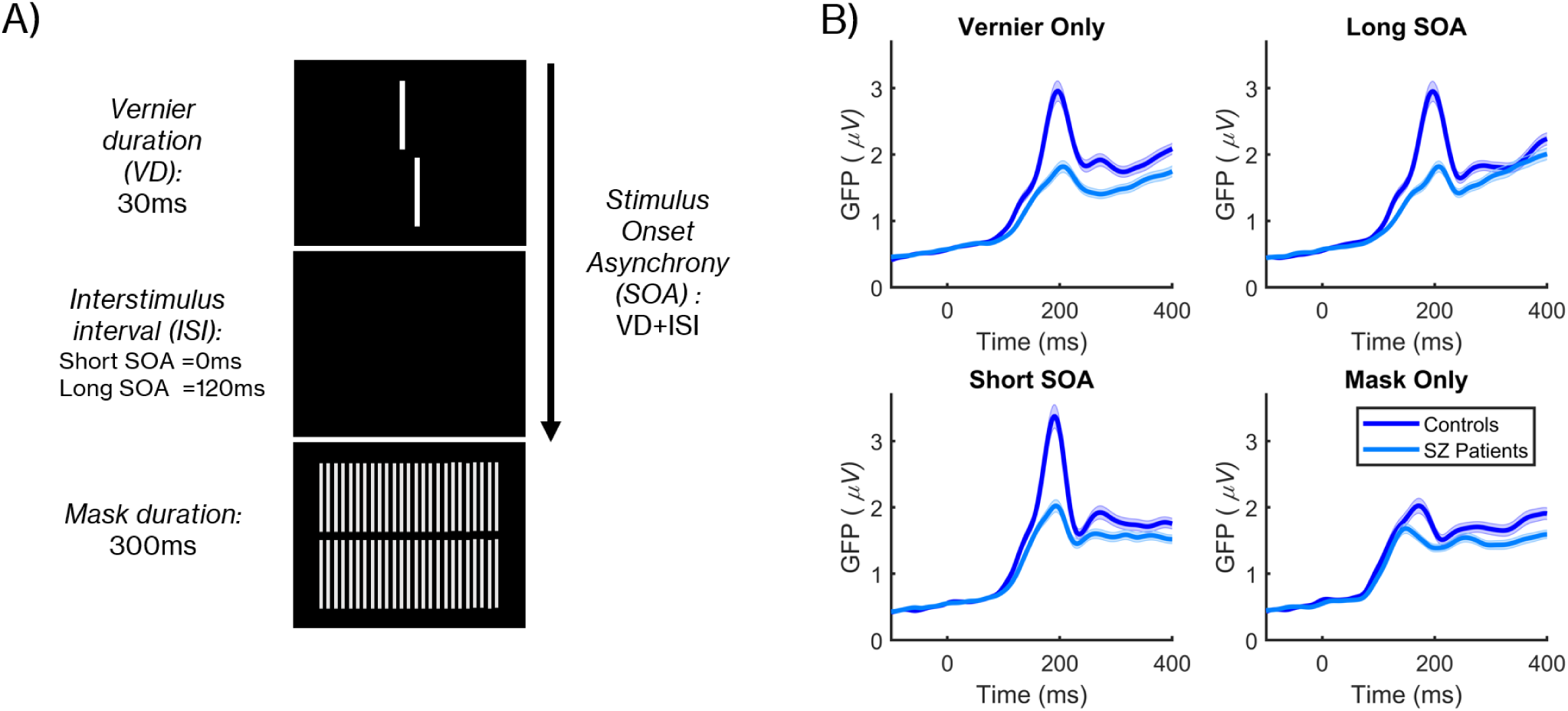
A) Shine-through visual backward masking (VBM) paradigm. Participants were presented with a Vernier (two vertical bars with a horizontal offset) followed by a grating mask^19^. Participants had to report the offset direction of the lower vertical bar relative to the upper one. In the Vernier Only condition, the Vernier was presented for 30ms. In the Short and Long SOA conditions, the Vernier was followed by a mask either with a stimulus onset asynchrony (SOA) of 30ms or 150ms. In the Mask Only condition, the mask was presented at the beginning of the trial for 300ms. B) Global field power (GFP) amplitudes for 142 patients with schizophrenia and 96 controls (data are pooled from previous studies^22,24,25^). Shaded error bars indicate the standard error of the mean (± sem).

**Figure 2.**
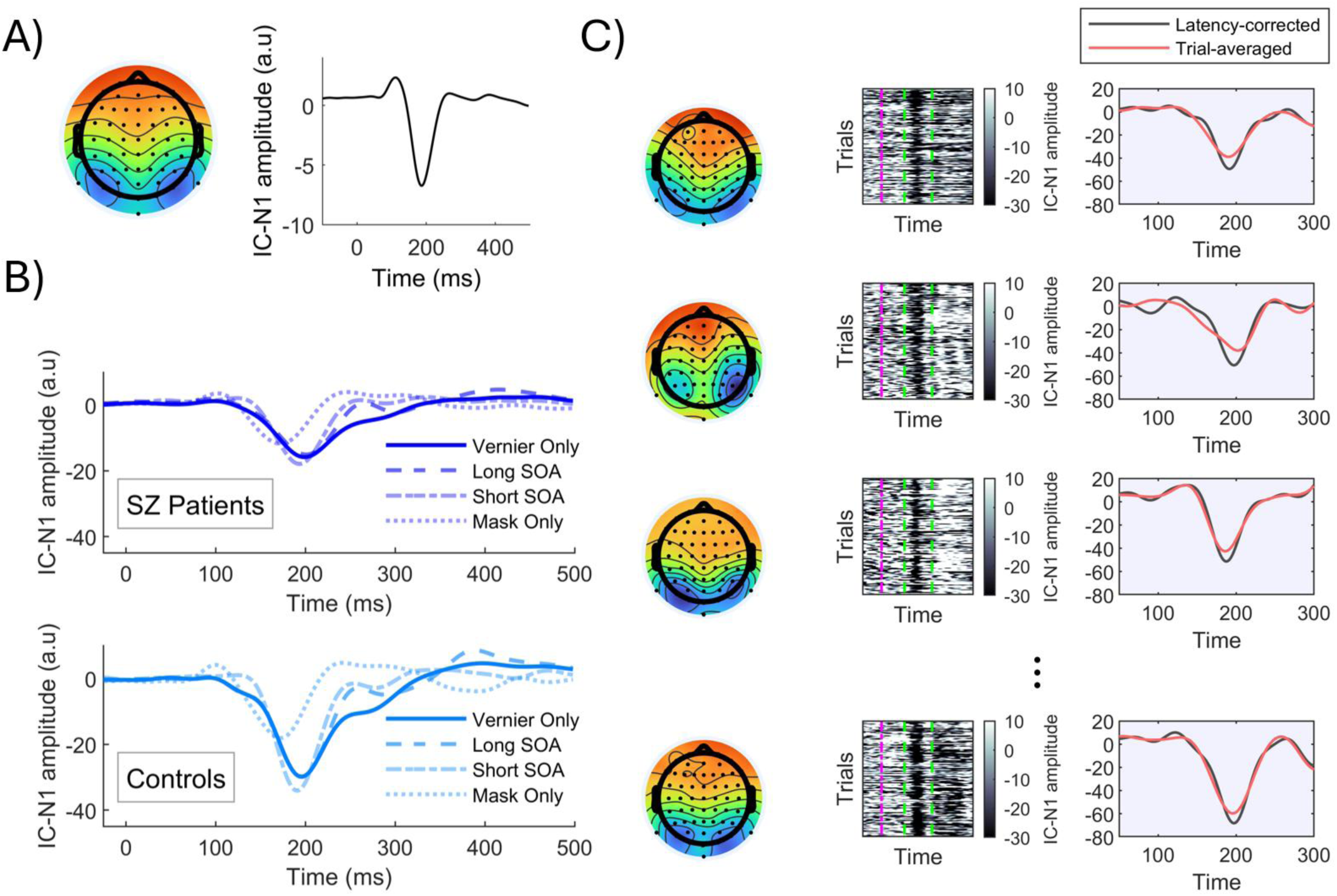
A) Group IC-N1 component obtained from the mlGICA procedure. B) IC-N1 component time courses for each group and condition. C) Example data from four participants. Left: individual-subject IC-N1 maps. Middle: single-trial plot of the IC-N1 response; purple dotted lines indicate stimulus onset, and green dotted lines indicate the window in which peaks were searched. Right: latency corrected N1 response and the corresponding trial-averaged waveform obtained without accounting for latency variability.

### IC-N1 amplitudes

Analysis of the IC-N1 amplitude before latency alignment revealed a significant main effect of CONDITION, F(3, 702)=91.208, P<0.001, 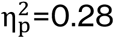 [0.226, 0.331], GROUP, F(1, 230)=25.727, P<0.001, 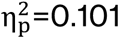 [0.039, 0.179], SEX, F(1, 230)=10.666, P=0.001, 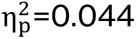 [0.007, 0.107], and a significant GROUP x CONDITION interaction, F(3, 702)=19.815, P<0.001, 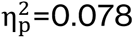 [0.042, 0.116]. No other main effect or interaction reached statistical significance (all P>0.05).

Post-hoc comparison of the GROUP x CONDITION interaction (Holm-adjusted) showed that while patients with schizophrenia exhibited significantly reduced IC-N1 amplitudes in all four conditions compared to controls, this difference was most pronounced in conditions with a target Vernier, with large effect sizes observed in the Vernier Only (P*corr*<0.001; d=-0.781 [-1.093, -0.468]), Long SOA (P*corr*<0.001; d=-0.821 [-1.133, -0.509]), and Short SOA (P*corr*<0.001; d=-0.927 [-1.239, -0.614]) conditions, and a small to medium effect size for the Mask Only condition (P*corr*=0.038; d=-0.331 [-0.644, -0.019]).

**Figure 3.**
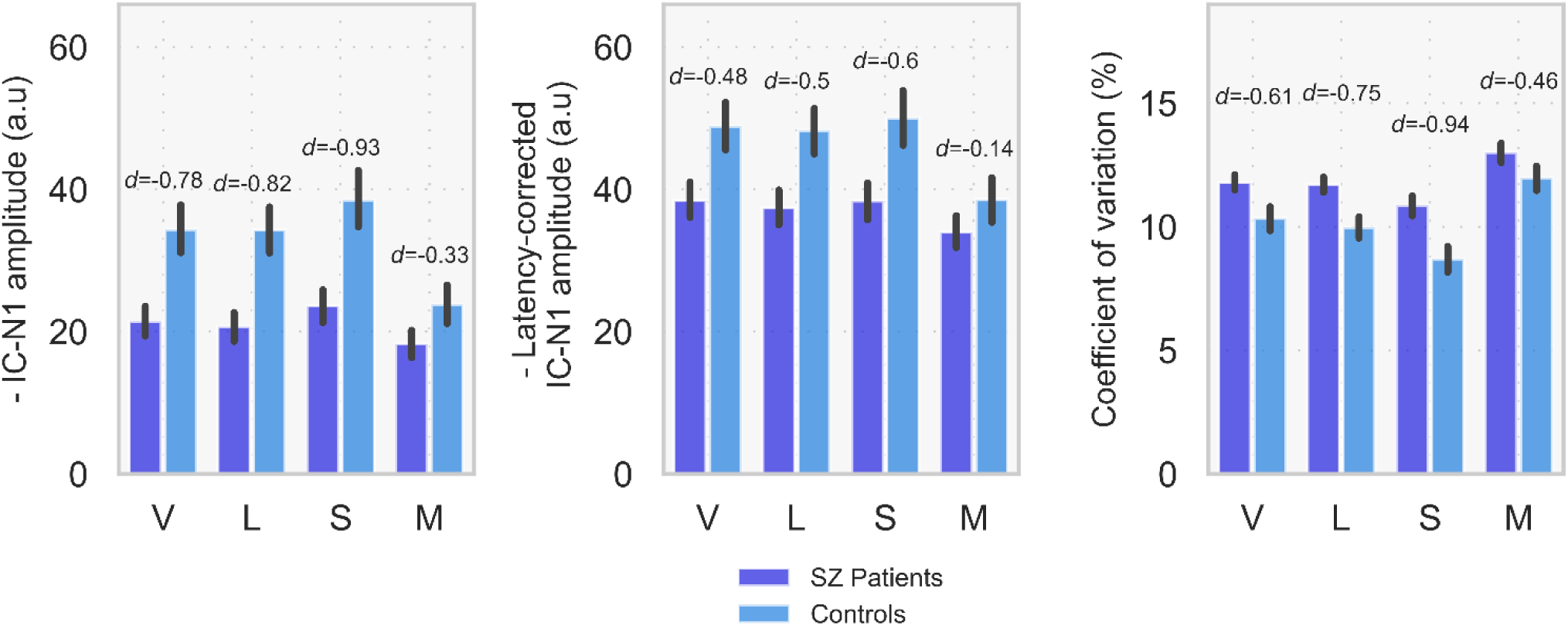
Left: IC-N1 amplitudes for each group and condition. Middle: latency-corrected IC-N1 amplitudes. Right: coefficient of variation (CV) calculated from the distribution of IC-N1 peak latencies; higher CV values indicate higher latency variability across trials. Positive values in the left and middle panels indicate higher amplitudes. V = Vernier Only, L = Long SOA, S = Short SOA, M = Mask Only. Error bars represent 95% confidence intervals obtained via bootstrapping. For each condition, Cohen’s d for the difference between patients and controls is shown above the corresponding bars.

Analysis of the latency-corrected IC-N1 amplitudes revealed a significant main effect of CONDITION, F(3, 702)=101.360, P<0.001, 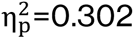 [0.248, 0.352], GROUP, F(1, 230)=10.311, P=0.002, 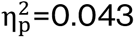 [0.006, 0.105], SEX(1, 230)=14.206, P<0.001, 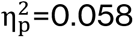 [0.014, 0.126], and a significant GROUP x CONDITION interaction, F(3, 702)=16.737, P<0.001, 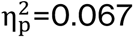 [0.033, 0.102].

Post-hoc analysis of the GROUP x CONDITION interaction revealed that patients showed significantly diminished amplitudes in all conditions except the Mask Only condition (P*corr*=0.369; d=-0.143[-0.457, 0.170]), compared to controls. The effect sizes were medium for all conditions with a target Vernier (Vernier Only: P*corr*=0.003; d=-0.484 [-0.798, -0.170], Long SOA: P*corr*=0.002; d=-0.5 [-0.814, -0.186]; Short SOA: P*corr*<0.001; d=-0.598 [-0.912, - 0.284]). No other main effects or interactions reached significance (P>0.05).

Importantly, the effect size for the group differences decreased after accounting for latency variability. For the Vernier Only condition, the effect size decreased from d=-0.781 to -0.484 (diff=0.297), in the Long SOA from d=-0.821 to -0.5 (diff=0.321), in the Short SOA condition from d=-0.927 to -0.598 (diff=0.329), and in the Mask Only condition from d=-0.331 to -0.143 (diff=0.188).

### IC-N1 coefficient of variation

We used the coefficient of variation (CV, %), as a measure of inter-trial variability. Analysis of the CV values revealed significant main effects of CONDITION, F(3, 702)=111.140, P<0.001, 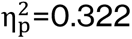 [0.268, 0.372], GROUP, F(1, 230)=30.891, P<0.001, 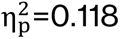 [0.051, 0.200], as well as a significant GROUP x CONDITION interaction F(3, 702)=4.297, P=0.005, 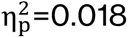 [0.002, 0.039].

Post-hoc analysis of the GROUP x CONDITION interaction revealed that patients with schizophrenia showed higher CV than controls in all conditions, yet with larger effect sizes for the conditions with the target Vernier. Effect sizes were large for the Long SOA (P*corr*<0.001; d=0.748 [0.442, 1.054]) and Short SOA conditions (P*corr*<0.001; d=0.940 [0.633, 1.246]), medium for the Vernier Only condition (P*corr*<0.001; d=0.609 [0.303, 0.915]), and small to medium for the Mask Only condition (P*corr*=0.003; d=0.463 [0.157, 0.769]).

To further characterize the GROUP × CONDITION interaction, we examined the simple effects of CONDITION within each group. Both controls and patients exhibited robust condition-dependent modulations in CV, with significantly higher CV in the Mask Only condition compared to all other conditions (all P*corr*<0.001), and lower CV in the Long SOA condition relative to Short SOA and Vernier Only conditions. This indicates that despite overall higher CV values, patients showed qualitatively similar context-dependent modulations of CV as controls.

We examined the relationship between CV and IC-N1 amplitudes (without latency correction) separately for each condition and group using Spearman correlations. All correlations were negative (i.e. higher CV values related to lower IC-N1 amplitudes) and remained significant after Holm correction for multiple comparisons within each group (all ρ = −0.47 to −0.65, P*corr*<0.001).

**Figure 4.**
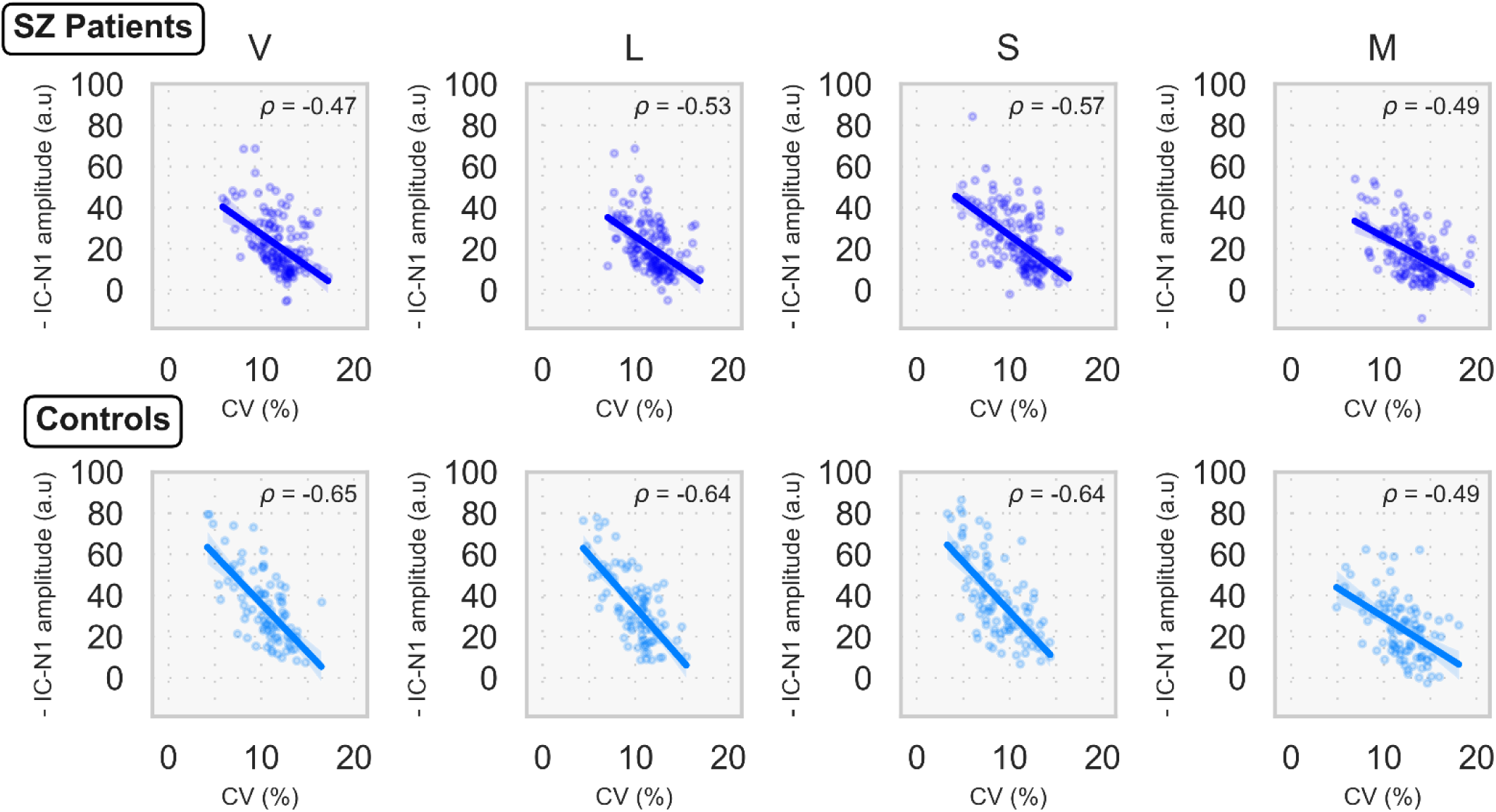
Scatter plots showing the relationship between the coefficient of variation (CV, %) and IC-N1 amplitudes for patients with schizophrenia and controls in the four conditions of the VBM experiment. Spearman correlation coefficients are shown for each plot. V = Vernier Only, L = Long SOA, S = Short SOA, M = Mask Only.

### CV associations to behavioral performance and medication

A beta regression mixed-effects model revealed significant main effects of CONDITION χ² (2)=1464.095, P<0.001, pseudo-R^2^=0.679, GROUP χ²(1)=46.108, P<0.001, pseudo-R^2^=0.062, and VISUAL ACUITY χ²(1)=12.707, P<0.001, pseudo-R^2^=0.018. Significant interactions were observed for CONDITION x GROUP χ²(2)=12.633, P=0.002, pseudo-R^2^=0.018, and CV × GROUP χ²(1)=6.378, P=0.012, pseudo-R2=0.009. All other effects were non-significant (all P>0.05).

Post-hoc analysis of CV slopes showed that CV was significantly predictive of behavioral performance in controls (P*corr*=0.002, slope=-0.244 [-0.389, -0.099]), whereas no significant relationship was observed in patients (P*corr*=0.968, slope=-0.002 [-0.105, 0.101]). This indicates that CV modulated accuracy in the control group only, despite both groups showing condition-dependent differences in performance. The effect sizes for this association were of medium size for the Vernier Only condition, and small for the Long SOA and Short SOA conditions in controls, as shown by Spearman rank correlations (Figure 5).

**Figure 5.**
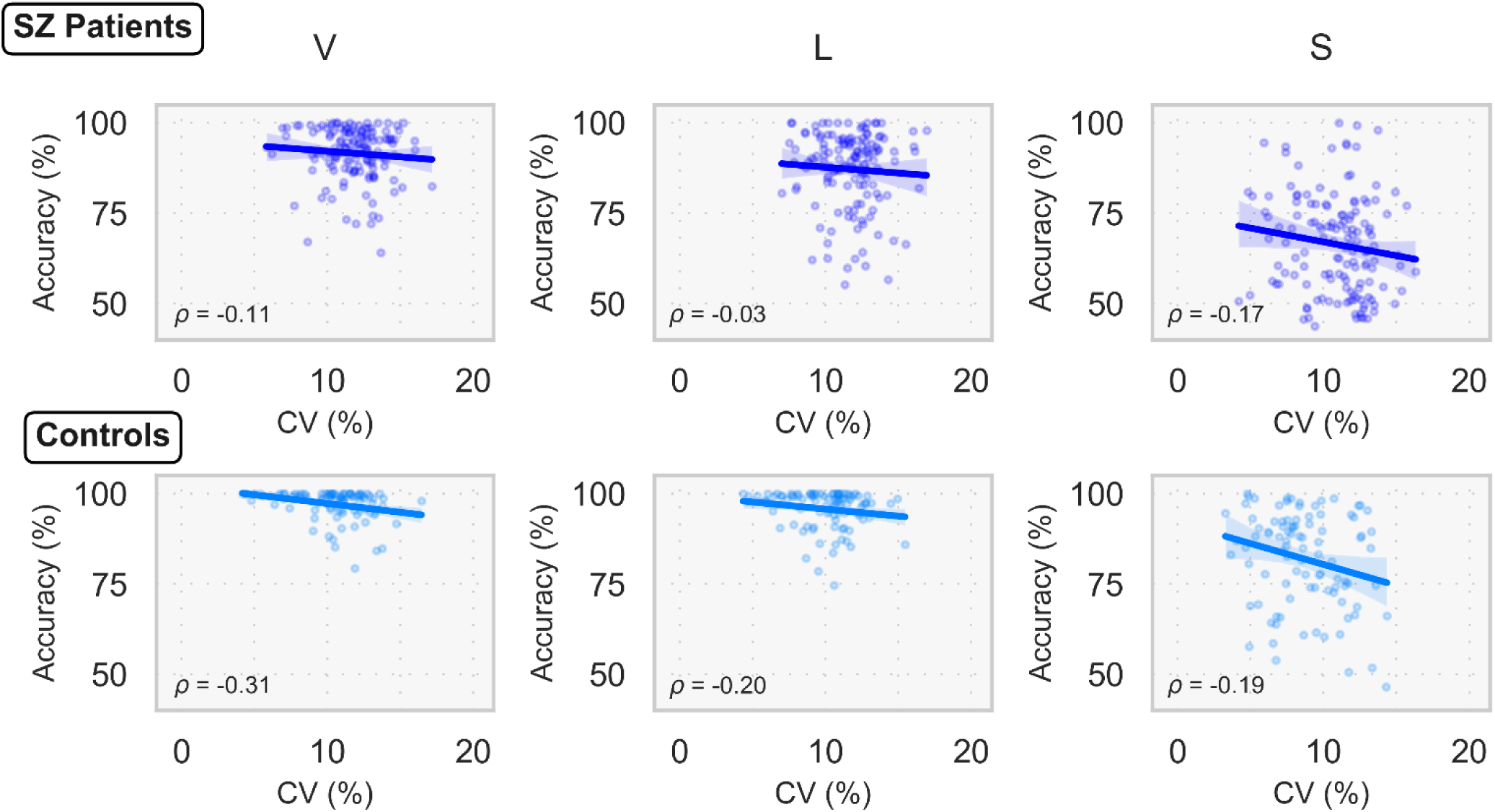
Scatter plots showing the relationship between the coefficient of variation (CV, %) and behavioral performance in the VBM task for controls and patients. Spearman correlation coefficients are shown for each plot. The Mask Only condition is not displayed because there were no correct responses. V = Vernier Only, L = Long SOA, S = Short SOA.

Analysis of the association between CV and chlorpromazine equivalents (CPZ) in patients revealed a significant main effect of CONDITION F(3, 378)=42.550, P<0.001, η_p_^2^=0.252 [0.179, 0.320] and a significant CONDITION × CPZ interaction F(3, 378)=5.578, P<0.001, η_p_^2^=0.042 [0.008, 0.083]. All other main effects, including symptom scores, and interactions did not reach statistical significance (all P>0.05).

To further characterize the CONDITION × CPZ interaction, we examined the simple slopes of CPZ predicting CV separately for each condition. CPZ significantly predicted CV in the Short SOA condition (P*corr*=0.035, slope=0.20 [0.051, 0.350]), whereas slopes for Vernier Only (P*corr*=0.304, slope=0.125 [–0.025, 0.274]), Long SOA (P*corr*=0.339, slope=0.105 [–0.045, 0.254]), and Mask Only (P*corr*=0.339, slope =–0.090 [–0.239, 0.060]) were not significant. Pairwise contrasts of CPZ slopes revealed that the Mask Only condition differed significantly from Vernier Only (P*corr*=0.020, Δslope=0.214 [0.018, 0.411]), Long SOA (P*corr*=0.036, Δslope=0.194 [-0.002, 0.390]), and Short SOA (P*corr*<0.001, Δslope=0.29 [0.094, 0.486]). All other slope differences between conditions were non-significant after Holm correction (all P*corr*>0.05).

## Discussion

In ERP studies, it is common to average trials to increase signal-to-noise ratio (SNR). Conclusions are drawn based on the average responses. However, if inter-trial variability is large, the average responses can be blurred and appear attenuated even if single-trial responses have high amplitudes, leading to biased results and conclusions. ERP studies in patients with schizophrenia have reported both reduced ERP amplitudes and higher inter-trial latency variability in several ERP components. Here, we examined to what extent latency variability contributes to reduced visual N1 responses in patients with schizophrenia.

We re-analyzed a large EEG dataset of 142 patients with schizophrenia and 96 healthy controls during a visual backward masking (VBM) experiment^22,24–26^. Previous studies with the same data demonstrated a strongly diminished N1 component in patients in comparison to controls. However, these studies have analyzed trial-averaged responses. Here, we used group ICA and identified an independent component resembling the visual N1 (i.e., IC-N1; Figure 2A) and estimated the latency of this component at the single-trial level (Figure 2C). We first found that the IC-N1 amplitudes mirrored the results from GFPs (Figure 1B, 2B), indicating significantly lower amplitudes in patients compared to controls, particularly pronounced in conditions with a target Vernier (Figure 3). Importantly, these group differences were still significant after accounting for latency variability in the analysis, although the effect sizes for the group differences decreased from large to medium, suggesting that reduced EEG responses in VBM can arise both from a weaker N1 response and from higher inter-trial latency variability in patients.

Our results are similar to those found for the P3b component, where the temporal consistency of the evoked responses across trials strongly predicted the trial-averaged amplitude^9^. This result was proposed to reflect an impairment of temporal imprecision of neuronal responses in patients, which might represent an important signature of schizophrenia. Our analysis of the variability of latency responses (CV) indicated that patients exhibited increased N1 latency variability in comparison to controls, in line with previous studies investigating other ERP components in patients^9–13^. However, although patients demonstrated increased CV values across conditions, the CV was modulated across experimental in a qualitatively similar manner than controls (Figure 3), indicating that increased latency variability might not reflect a general deficit of temporal imprecision, but an inability to adjust neural variability according to the task demand to the same extent as controls.

To understand the contribution of neural variability to behavioral deficits in patients, we examined the relationship between CV to VBM behavioral accuracy, since it has previously been proposed that reduced temporal consistency of neural responses can result in reduced behavioral performance^6,27^. We found a significant relationship between CV and behavioral performance yet only in controls, suggesting a weaker or absent contribution of CV to behavior in patients. Our analysis of the relationship between CV and CPZ provides evidence that antipsychotic medication is a contributing factor. The relationship between CV and CPZ, although modest in effect size, was most strongly pronounced in conditions with a target vernier, and not in the Mask Only condition, indicating that higher CPZ values might affect neural responses only when a task-relevant element needs to be processed. This again highlights that temporally imprecise neural responses arise more strongly in patients when task demands are higher.

Impairments in recurrent processing have been proposed to impede patients from enhancing and stabilizing visual information, and therefore they could explain reduced amplitudes and higher inter-trial variability in VBM^18^. One candidate mechanism for recurrent processing is attention, which can modulate neural responses in sensory areas both by boosting neural activity^28,29^, and also by reducing the variability of neural responses across trials^30^, and which is strongly impaired in schizophrenia^31^. Importantly, we found that in the Mask Only condition, patients and controls showed similar IC-N1 amplitudes, suggesting that diminished responses arise more strongly when there is a visual target and higher attention and neural resources are required This is in line with previous studies showing that VBM impairments in schizophrenia probe a deficit in target enhancement, i.e., a top-down deficit, instead of a deficit in early visual processing^18^.

In particular, we proposed that the cholinergic system stabilizes the faint information of the briefly presented vernier to protect it from the mask and that this system is impaired in the patients-either being reduced in amplitude or less precise in time^18^. Our results show that both aspects play a role. For instance, while latency variability strongly correlated with IC-N1 amplitudes, the explained variance was on average 27% in patients, indicating that, in some patients, reduced peak amplitudes cannot be explained by increased latency variability. Cholinergic input has been shown to stabilize attractor dynamics in prefrontal cortex and reduce internal variability^32^. Similar principles may also apply to sensory circuits. Given that antipsychotics often have anticholinergic properties, the observed relationship between CV and CPZ may suggest that EEG-derived inter-trial variability might, to some degree, be sensitive to cholinergic modulation. Future studies will be important to clarify the role of the cholinergic system in shaping neural variability in visual cortex when faint information needs to be enhanced. Moreover, future studies using anticholinergic burden scales instead of CPZ might help clarify this relationship. The duration of the N1 response has also been previously suggested to be a neural correlate for target enhancement^33^. Hence, evaluating these and other features of the ERPs in a *multiverse* study could inform on the shared mechanisms across features^34,35^.

Our results argue for a reinterpretation of reduced ERP responses in patients with schizophrenia. As we demonstrate, latency variability is a contributing factor to reduced ERPs. Hence, reduced ERPs might not necessarily reflect weaker activations only. Further advances in the study of temporal variability may help identify the impaired underlying neural mechanisms and how they contribute to visual processing dysfunctions in patients. Because increased latency variability has been reported across several ERP components in schizophrenia, a common dysfunction of temporal variability could be a contributing factor to deficits observed across paradigms. This hypothesis remains to be tested systematically using batteries of EEG paradigms.

## Methods

### Participants

Data from 142 patients with schizophrenia (including from 19 first-episode psychosis patients previously published in ^26^) and 96 healthy controls were analyzed. All data except from two healthy controls and four patients with schizophrenia have been published^24–26^. Patients were recruited from the Tbilisi Mental Health Hospital, the psycho-social rehabilitation center, or the acute psychiatric departments of the multi-profile clinics. The Diagnostic and Statistical Manual of Mental Disorders Fourth Edition (DSM-IV) was used for clinical diagnoses through a Structured Clinical Interview. The severity of psychopathology was assessed using the Scale for the Assessment of Positive Symptoms (SAPS) and the Scale for the Assessment of Negative Symptoms (SANS). Healthy controls with no family history of psychosis were recruited from the general population of Tbilisi, Georgia. Participant age ranged from 16 to 55 years old. General exclusion criteria were substance abuse and the presence of severe developmental, neurological, or somatic disorders. All procedures received approval from the local ethics committee and complied with the Declaration of Helsinki. All participants provided written informed consent and were informed that they could quit the experiment at any time.

**Table 1.**
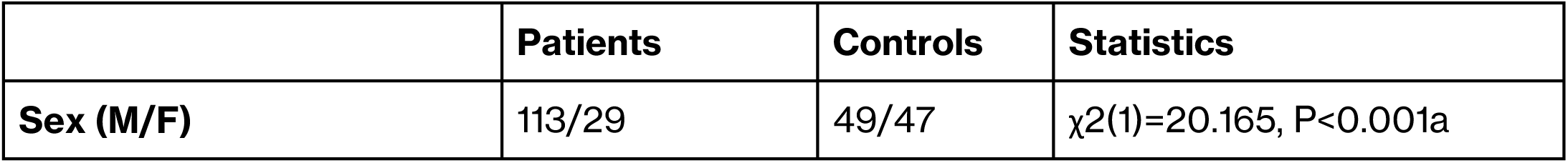
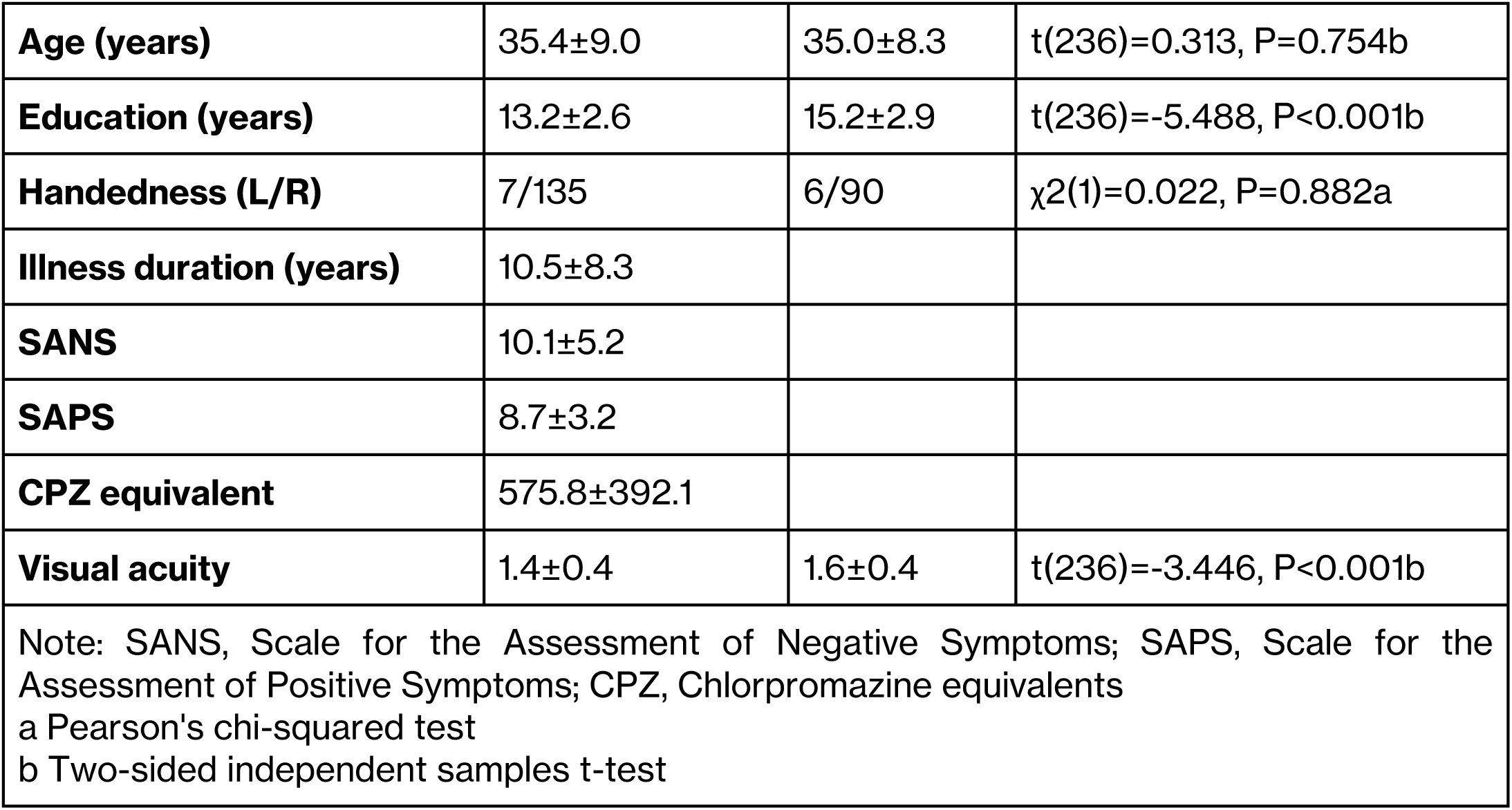
Group average statistics of patients with schizophrenia and controls.

### Experiment

*Apparatus.* Visual stimuli were presented on a cathode ray tube (CRT) Siemens Fujitsu P796-1 screen, which had a refresh rate of 100 Hz and a screen resolution of 1024 x 768 pixels. Participants were seated 3.5m from the screen in a dimly lit room. Each pixel on the screen occupied an angle equivalent to about 0.3’ (arcminutes). The stimuli appeared in white against a black background with a luminance of 100 cd/m². The black background had a luminance of < 1 cd/m².

*Stimuli.* The vernier stimulus consisted of two vertical line segments of 10’ length with a gap of 1’. The lower segment was slightly shifted to the left or right relative to the upper one, with a fixed offset of approximately 1.2’. The mask was composed of 25 verniers without offset separated by 3.33’.

*EEG experiment.* The vernier duration (VD) was set to 30 milliseconds which corresponds to the average VD observed in previous studies^17,19^. The experiment comprised four stimulus conditions (Vernier Only, Short SOA, Long SOA, and Mask Only^22,26,36^). In the Vernier Only condition, only the target vernier was presented. In the Long SOA and Short SOA conditions, a mask following the target vernier was presented with a stimulus onset asynchrony (SOA) of 150 or 30 milliseconds, respectively. The choice of SOAs for the different conditions was informed by the mean SOAs observed in previous studies involving both patients and controls^17,19,22,26^. In the Mask Only condition, the mask stimulus was presented at the beginning of the trial for 300ms. Participants had to report the perceived offset direction of the lower bar (left or right) compared to the upper bar of the vernier stimulus (Figure 1A).

### EEG recording and preprocessing

EEG was recorded with an ActiveTwo Mk2 system (BioSemi B.V., The Netherlands) using 64 Ag-AgCl sintered active electrodes, referenced to the common mode sense electrode, with a sampling rate of 2048 Hz. Data were down-sampled to 512 Hz and preprocessed using an automatic pipeline (APP^37^). Preprocessing included: band-pass filtering between 1 and 40 Hz; re-referencing to the bi-weight estimate of the average of all electrodes; removal and 3D spline interpolation of bad electrodes; removal of bad epochs; independent component analysis (ICA) to attenuate ocular and muscle activity, and removal of epochs with artifacts. Data were then baseline corrected (0.5s baseline) and re-referenced to the average of all channels. Trials containing values exceeding ±90uV were rejected. EEG data were further low-pass filtered at 20Hz before ERP analyses. Data from four patients with schizophrenia and one control were discarded due to excessive muscular artifacts, bad electrodes, or artifact trials.

### Topographic Segmentation Analysis

We used topographic segmentation analysis^38^ as an initial step to find the N1 component map, which was used later to identify the independent component that most strongly resembled the N1 response. A grand average EEG dataset (averaged across trials, conditions, subjects, and groups) was subjected to a *k*-means clustering algorithm. The *k*-means procedure finds distinct topographic maps that are present at different time points and explain most of the variance of the dataset. The *k* value (i.e., the number of topographic maps) was set to 5 as a trade-off between the number of maps and interpretability. We used the Microstate-EEGlab toolbox^39^.

To identify the topographic map corresponding to the N1 component, we backfitted the 5 maps to the average EEG datasets of each group and condition (i.e., to 8 datasets, 4 for controls and 4 for patients). We then identified the map that was present at the GFP peak of these datasets (Figure S1).

### Independent component analysis

Multi-level group independent component analysis (mlGICA) was used to isolate brain patterns of activity (i.e., independent components) that are present across datasets^40,41^. The main advantage of this approach is that isolating the components leads to an increase in SNR, which facilitates single-trial EEG analyses^42^.

In mlGICA, two data reduction steps are applied before ICA. Principal component analysis (PCA) is commonly used before ICA to reduce the complexity of the analysis by minimizing dependencies between components^41^. Moreover, typically around 10-20 components explain most of the variance in the EEG data. We followed this procedure and reduced the data of each participant using PCA, retaining the first 10 principal components which on average explained 77% of the variance. To obtain the ICA solution, we used the first 100 trials of each dataset since the number of trials had to be the same across participants for the next step of the analysis. Data from all participants were vertically concatenated into a new matrix. The resulting matrix of size (NSubjects x NComponents) x (NTrials x NTimePoints x NConditions) was then subjected to a second PCA, from which the first 10 principal components were retained. ICA was applied to these 10 components. Each of the 10 independent components (ICs) was correlated with the N1 map obtained from the topographic segmentation analysis, and the IC with the highest correlation was defined as the IC-N1 map (Figure S1).

For single-trial analysis, the time courses and topographic maps of the IC-N1 were back-reconstructed for each participant individually using the PCA coefficients and the ICA demixing matrix. IC-N1 time courses were back-reconstructed using all available trials (across all four conditions; patients: M=574.87, SD=33.35; controls: M=582.41, SD=29.96).

### Single-trial latency analysis

To obtain the latency of the IC-N1 for each trial, we searched for the time point at which the amplitude reached its minimum value (i.e., the negative peak). We selected a 100ms window (within 125-275ms post-stimulus) for peak detection based on our previous results^22,24,25^. This window was centered on the time at which the trial-averaged IC-N1 reached its peak amplitude. For example, if for one participant the trial-averaged peak amplitude occurred at 200ms, the time window to search for the single-trial peaks was 150-250ms. From the distribution of peak latencies, we removed the peaks that were found at the edges of the analysis window. Further outliers were detected using 1.5 IQR bounds to discard potentially unreliable peaks. We calculated the coefficient of variation (CV), dividing the standard deviation by the mean peak latency and used CV as the measure for inter-trial variability.

To estimate the latency-corrected IC-N1 amplitude, we averaged the single-trial peak values, irrespective of when each peak occurred within the specified time window. We also applied two other algorithms to estimate peak latencies. The first was a template-matching procedure^43^, in which the average response across trials is first computed and used as a template; single trials are then shifted in time until the correlation to the template is maximized. We used a single iteration of this algorithm (i.e., the template was computed only once) to obtain the latency corrected response. In the second method, first, a dimensionality reduction algorithm based on Laplacian embedding was first applied to the EEG data to find a relevant axis of variation, representing latency variability across trials. Importantly, isolating the components using ICA improves the performance of this procedure, which assumes that a single parameter (the response latency) changes across trials. Finally, the latency of single trials was calculated using a graph-cuts-based approach^42^.

### Statistical analysis

Statistical analyses were performed using R (version 4.5.1). For statistical analysis of the IC-N1 amplitudes, latency-corrected IC-N1 amplitudes, CV, and the relationship between clinical variables and CV, we used linear mixed-effects models fitted with the lme4 (version 1.1-37) package.

The general model structure was:

- *Variable (CV/IC-N1/Latency-corrected IC-N1) ∼ GROUP:VISUAL ACUITY + GROUP:EDUCATION + SEX*GROUP*CONDITION + (1|ID)*

- *CV ∼ VISUAL ACUITY + ILLNESS DURATION + EDUCATION + SANS + SAPS + CONDITION*CPZ + (1|ID)*

Fixed-effects significance was assessed using Type II Wald F-tests (via the car 3.1-3 package). Effect sizes were quantified as partial eta squared (η²) with 95% confidence intervals, computed using the effectsize 1.0.1 package.

For the analysis of behavioral accuracy, we fitted generalized linear mixed models using the glmmTMB 1.1.9 package. Because the dependent variable (accuracy) was bounded between 0 and 1, we modeled it with a beta distribution and a logit link function. To account for condition-specific variability, the dispersion parameter was modeled as a function of CONDITION. Fixed-effects significance was assessed using Type II Wald χ² tests. All post-hoc analysis P-values were Holm-adjusted and calculated using the emmeans 1.11.2 package.

- *Accuracy ∼ VISUAL ACUITY + EDUCATION + SEX + LATENCY-CORRECTED IC-N1 + CV*CONDITION*GROUP + (1|ID)*

## Author Contributions

D.G. and J.R.C. conducted the analysis and drafted the manuscript. M.H.H., E.C., A.B., and P.F. supervised the research and contributed to manuscript editing. M.R. collected the data and contributed to manuscript editing.

