## Supplementary Figures for "Higher inter-trial latency variability contributes to reduced visual EEG responses in schizophrenia"

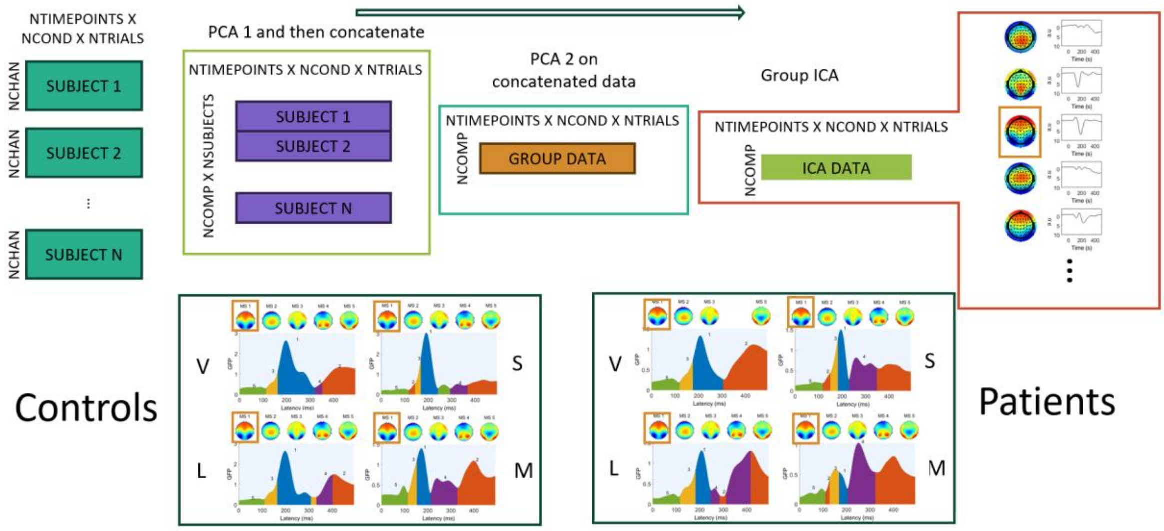


Figure S1. Illustration of group ICA procedure, and topographic segmentation analysis. Before ICA, two PCAs were applied to find common components across datasets. The resulting ICs were then compared to the maps obtained using topographical segmentation analysis (lower panel). We found that maps obtained from topographic segmentation analysis, which were active around 200ms, strongly resembled one of the independent components. We defined that independent component as the IC-N1 component. V=Vernier Only, L=Long SOA, S=Short SOA, M=Mask Only.

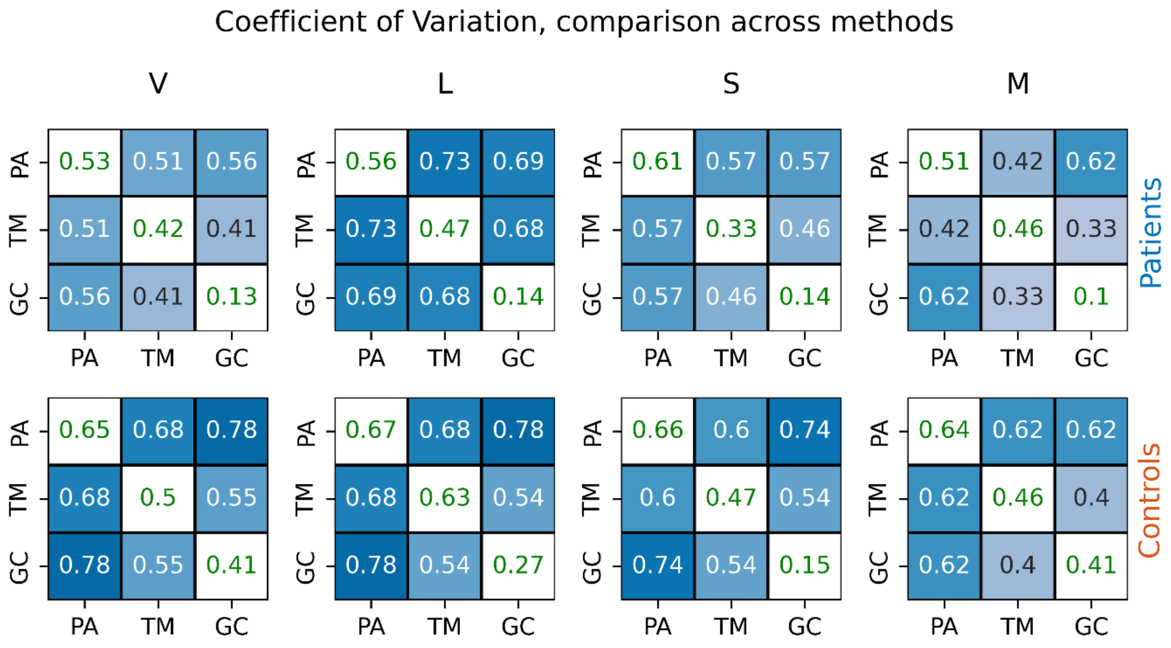


Figure S2. Correlation between the outputs of the different methods to find peak latencies. PA: Using the minimum value within a pre-defined window (reported in the main text); TM: Template matching; GC: Method based on graph cuts. The values in green fonts presented in the diagonals indicate the within session stability of the method. Stability was calculated using intraclass correlations between using the CV obtained from the first and last 50 trials for all methods. V=Vernier Only, L=Long SOA, S=Short SOA, M=Mask Only.
